# High-Throughput Sequencing of Transposable Element Insertions Suggests Adaptive Evolution of the Invasive Asian Tiger Mosquito Towards Temperate Environments

**DOI:** 10.1101/049197

**Authors:** Clément Goubert, Hélène Henri, Guillaume Minard, Claire Valiente Moro, Patrick Mavingui, Cristina Vieira, Matthieu Boulesteix

**Affiliations:** Université de Lyon, F-69622, Lyon, France; Université Claude Bernard Lyon 1, CNRS, Laboratoire de Biométrie et Biologie Evolutive, UMR5558, F-69100 Villeurbanne, France; Université de Lyon, F-69622, Lyon, France; Université Lyon 1, Villeurbanne, France; CNRS, UMR 5557, Ecologie Microbienne, Villeurbanne, France; INRA, UMR 1418, Villeurbanne, France; Metapopulation Research Center, Department of Biosciences, University of Helsinki, Helsinki, Finland; Université de La Réunion, UMR PIMIT, INSERM 1187, CNRS 9192, IRD 249, Plateforme Technologique CYROI, Sainte-Clotilde, La Réunion.

**Keywords:** Invasive species, *Aedes albopictus*, Local Adaptation, Genome Scan, Transposable Elements

## Abstract

Invasive species represent unique opportunities to evaluate the role of local adaptation during colonization of new environments. Among these species, the Asian tiger mosquito, *Aedes albopictus*, is a threatening vector of several human viral diseases, including dengue and chikungunya, and raises concerns about the Zika fever. Its broad presence in both temperate and tropical environments has been considered the reflection of great “ecological plasticity”. However, no study has been conducted to assess the role of adaptive evolution in the ecological success of *Ae. albopictus* at the molecular level. In the present study, we performed a genomic scan to search for potential signatures of selection leading to local adaptation in one-hundred-forty field-collected mosquitoes from native populations of Vietnam and temperate invasive populations of Europe. High-throughput genotyping of transposable element insertions led to the discovery of more than 120 000 polymorphic loci, which, in their great majority, revealed a virtual absence of structure between the bio-geographic areas. Nevertheless, 92 outlier loci showed a high level of differentiation between temperate and tropical populations. The majority of these loci segregates at high insertion frequencies among European populations, indicating that this pattern could have been caused by recent adaptive evolution events in temperate areas. An analysis of the overlapping and neighboring genes highlighted several candidates, including diapause, lipid and juvenile hormone pathways.

## Introduction

Biological invasions represent unique opportunities to study rapid evolutionary changes, such as adaptive evolution. Settlement in a novel area is a biological challenge that invasive species have successfully overcome. The underlying processes can be studied at the molecular level, particularly to gather empirical knowledge of the genetics of invasions, a field of study that has produced extensive theoretical predictions for which there is still little evidence in nature (Colautti & Lau 2015). Some of the main concerns are disentangling the effects of neutral processes during colonization, such as founder events or allele surfing at the migration front, from adaptive evolution (i.e., local adaptation, Lande 2015; Peischl & Excoffier 2015; Colautti & Lau 2015).

Signatures of adaptation can be tracked on genomes due to characteristic patterns of reduced genetic diversity left by the appearance and spread of a new beneficial mutation (Tajima 1989; Braverman *et al*. 1995; Fay & Wu 2000; Nielsen 2005; Vitti *et al*. 2013). Contrasting regimes of selection between populations can also leave high levels of genetic differentiation in the vicinity of adaptive loci (Lewontin & Krakauer 1973; Maynard Smith & Haigh 1974). The strength of such signals can be influenced by the origin of the adaptive mutation, e.g., if it arises *de novo* or if it spreads from the standing genetic variation (Pritchard *et al*. 2010; Messer & Petrov 2013). However, detection of the footprint of natural selection is dependent on the availability of informative genetic markers, which should ideally provide substantial coverage of the genome to allow selection scans and be easily and confidently scored across many individuals.

Unfortunately, invasive organisms are rarely model species, making the development of a reliable and efficient marker challenging.

The Asian tiger mosquito, *Aedes* (*Stegomya*) *albopictus* (Diptera:Culicidae) is currently one of the most threatening invasive species (Invasive Species Specialist Group). Originating from Southeastern Asia, it is one of the primary vectors of dengue and chikungunya viruses and is also involved in the transmission of other threatening arboviruses (Paupy *et al*. 2009), in particular, the newly emerging Zika virus (Grard *et al*. 2014; Marcondes & Ximenes 2015; Chouin-Carneiro *et al*. 2016). Nowadays, *Ae. albopictus* has settled in every continent except Antarctica and is found in both tropical and temperate climates (Bonizzoni *et al*. 2013). Though this species is supposed to have emerged from rain forests (Hawley 1988), the acknowledged native area of *Ae. albopictus* encompasses contrasting environments including temperate regions of Japan and China, offering a large potential of fit towards the most recently colonized environments. For example, the induction of photoperiodic diapause in temperate areas, which has a genetic basis in *Ae. albopictus* (Hawley *et al*. 1987; Hanson & Craig 1994; Urbanski *et al*. 2010), is decisive to ensure invasive success in Europe or Northern America. It allows the susceptible populations to survive through winter at the larval stage into eggs. Such a trait appears to be governed by a “genetic toolkit” involving numerous genes and metabolic networks for which the genetic polymorphism between diapausing and non-diapausing strains remains to be elucidated (Poelchau *et al*. 2013a). In addition, the colonization of new areas that appear similar to the native environment at first glance can still involve *de novo* adaptation; even environments that share climatic variables are not necessarily similar regarding edaphic and biotic interactions (Colautti & Lau 2015). This suggests that, regardless of the native and settled environments, it might be possible to find evidence of adaptive evolution in invasive populations of *Ae. albopictus*.

To better understand the invasive success of this species, we genotyped 140 field individuals collected from three Vietnamese (native tropical area) and five European (invasive temperate area) populations (Figure 1), aiming to identify genomic regions involved in local adaptation. To do so, we developed new genetic markers based on high-throughput genotyping of the insertion of transposable elements (TEs) that represent at least one third of the genome of *Ae. albopictus* and include recently active families that can reach thousands of copies in one genome (Goubert *et al*. 2015).

**Figure 1.**
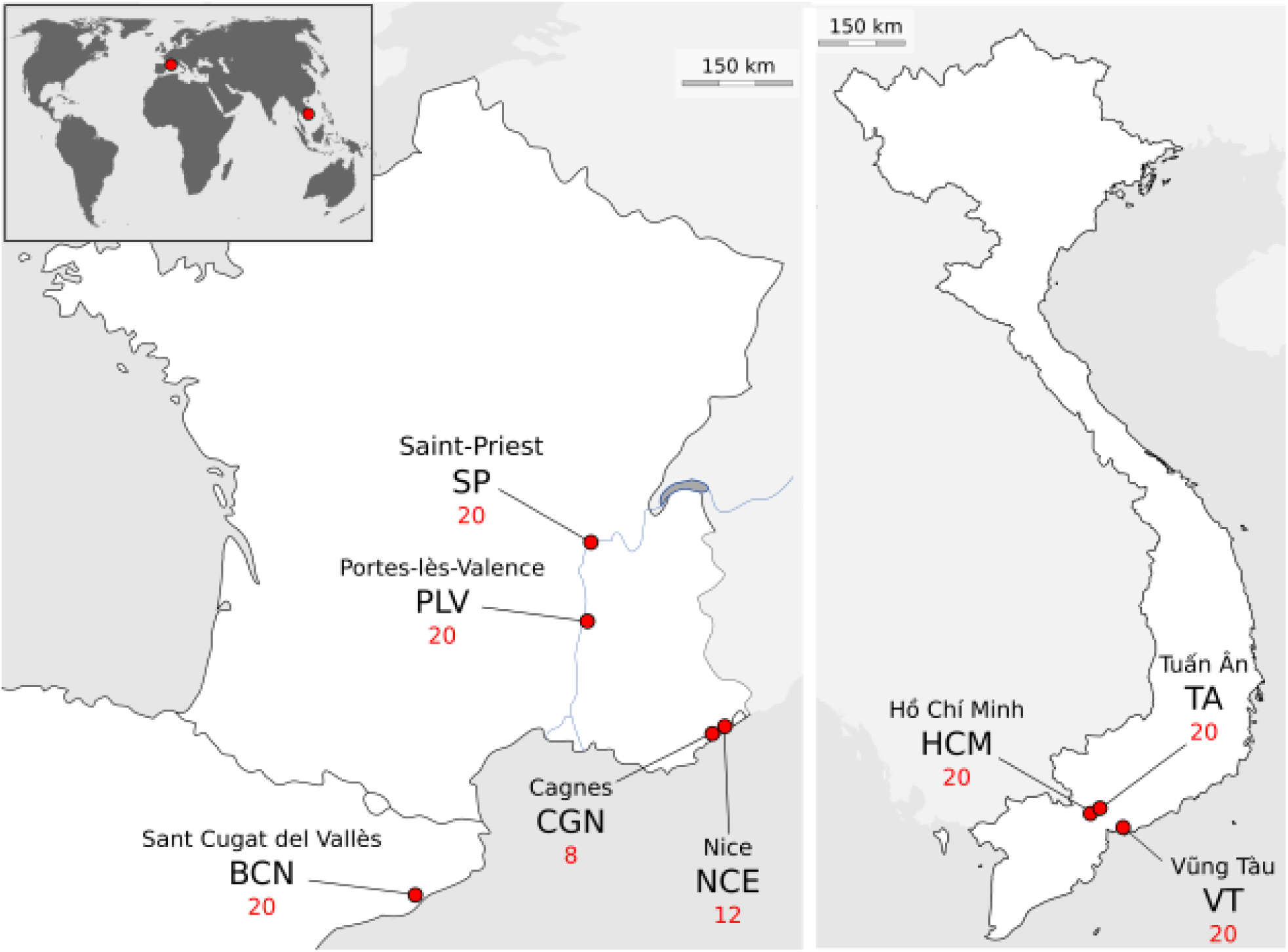
Sampling sites (with abbreviations) of *Ae. albopictus* in Europe and Vietnam. Red numbers correspond to the total number of individuals sampled. Supporting information on samples is available in Table S1.

Amplification of TE insertions is particularly efficient to obtain many genetic markers throughout one genome (Bonin *et al*. 2008), especially if few genomic resources are available (Monden *et al*. 2014), which was the case for the Asian tiger mosquito until recently. In addition, such markers represent an attractive alternative to other methods of diversity reduction such as RAD-sequencing (Miller et al. 2007) that could be less efficient in species with a high TE load (Davey et al. 2012) and did not produce satisfying results in *Ae. albopictus* (Goubert *et al*. 2016). In mosquitoes, TEs have been shown to be powerful markers for both population structure analysis (Biedler *et al*. 2003; Boulesteix *et al*. 2007; Santolamazza *et al*. 2008; Esnault *et al*. 2008) and genome scans (Bonin *et al*. 2009).

We hypothesized that some TE insertion sites could be located in the neighborhood of targets of natural selection and thus could reach a high level of differentiation between native and invasive populations if selective sweeps occurred during local adaptation. In addition, some TEs could also insert near or inside coding regions and many studies have shown their recurrent involvement in environmental adaptation in multiple organisms (Casacuberta & González 2013), eventually contributing to the success of invasive species (Schrader et al. 2014, Stapley et al. 2015).

To distinguish between neutral demographic effects and adaptive evolution, we first performed population genetic analyses to reveal the global genetic structure of the studied populations. We then performed a genomic scan for selection and identified 92 candidate loci under directional selection, among which several are located within or in a close neighborhood of annotated genes, revealing candidate pathways to investigate in forthcoming studies.

## Materials and Methods

### Biological samples

One hundred-forty flying adult female *Ae. albopictus* were collected in the field at eight sampling sites in Europe and Vietnam during the summers of 2012 and 2013 (Figure 1 and Table S1). Individuals were sampled using either a single trap or aspirators through the sampling site within a 50-meter radius. When traps were used, live mosquitoes were collected after a maximum of two days.

### High-throughput transposon display (TD) genotyping

The insertion polymorphism of the five transposable element families (I Loner Ele1 (IL1), Loa Ele2B (L2B), RTE4, RTE5 and Lian 1) identified by Goubert *et al*. (2015) in *Ae. albopictus* was characterized using transposon display (TD), a TE insertion-specific PCR method, combined with Illumina sequencing of all TD amplification products (Figure S1). These TE families were chosen according to high copy number estimate (from 513 to 4203 copies), high identity between copies, and a “copy and paste” mode of transposition (all these TEs are non-LTR retrotransposons).

#### DNA extraction and TD adapter ligation

The total DNA was extracted from whole adult bodies following the phenol-chloroform protocol described by Minard *et al*. (2015). The TD was conducted combining methods from previous studies (Munroe *et al*. 1994; Roy *et al*. 1999; Akkouche *et al*. 2012; Carnelossi *et al*. 2014). First, individual extracted DNA (≈ 75 ng) was used for enzymatic digestion in a total volume of 20 *μ*L, with HindIII enzyme (10 U/*μ*L) and buffer R (Thermo Scientific) for 3 hours at 37°C. The enzyme was inactivated at 80°C for 20 minutes. TD adapters were then built by hybridizing Hindlink with MSEB oligonucleotides (100 *μ*M, see Table S2) in 20× SSC and 1 M Tris in a total volume of 333 *μ*L after 5 min of initial denaturation at 92° C and 1 h at room temperature for hybridization. Once ready, the TD adapters were ligated to 20 *μ*L of the digested DNA by mixing 2 *μ*L of TD adapter with 10 U T4 ligase and 5× buffer (Fermentas) in a final volume of 50 *μ*L for 3 hours at 23°C.

#### Library construction

For each individual and for each of the five TE families, the TE insertions were amplified by PCR (PCR 1) in a Biorad Thermal Cycler (C1000 or S1000) in a final volume of 25 *μ*L The mixture contained 2 *μ*L of digested-ligated DNA with 1 *μ*L dNTPs (10 mM), 0.5 *μ*L TD adapter-specific primer (LNP, 10 *μ*M, see S3 Table) and 0.5 *μ*L of TE-specific primer (10 *μ*M), 1 U AccuTaq polymerase (5 U/*μ*L) with 10× buffer and dimethyl sulfoxide (Sigma). Amplification was performed as follows: denaturation at 98°C for 30 seconds then 30 cycles of 94°C for 15 seconds, hybridization at 60°C for 20 seconds and elongation at 68°C for 1 minute; a final elongation was performed for 5 minutes at 68°C. For L2B and RTE5 TEs, a nested PCR was performed to increase specificity under the same PCR conditions using internal forward TE primers and LNP (Table S2). The PCR 1 primers include a shared tag sequence that was used for hybridization of the individual indexes by PCR 2.

Multiple independent PCR1 can be performed to avoid amplification bias during the library preparation (Recknagel *et al*. 2015). Accordingly, three independent PCR 1 were performed from the same digestion product for each TE family. The PCR 1 products (3 PCR * 5 TE per individual) were then purified using volume-to-volume Agencout AMPure XP beads (20 *μ*L PCR 1 + 20 *μ*L beads) and eluted in 30 *μ*L resuspension buffer. After NanoDrop quantification, equimolar pools containing the 3*5 PCR products per individual were made using a Tecan EVO200 robot. Individual pools were then size selected for 300 to 600 bp fragments using Agencout AMPure XP beads as follows: first, the magnetic beads were diluted in H2O at a 1:0.68 ratio then added to 0.625 × PCR products to exclude long fragments. A second purification was performed using a non-diluted bead:DNA ratio of 1:8.3 to exclude small fragments.

Multiplexing samples was performed using a homemade 6 bp index (included in SRA individual name), which was added to the R primer (Table S2) during a second PCR (PCR 2) with 12 cycles in an ABI 2720 Thermal Cycler. The mixture contained 15 ng PCR products, 1 *μ*l of dNTPs (10 mM), 0.5 *μ*l MTP Taq DNA polymerase (5 U/*μ*l, Sigma), 5 *μ*l 10X MTP Taq buffer and 1.25 *μ*l of each tagged primer (20 *μ*m) in a final volume of 50 *μ*l. Amplification was performed as follows: denaturation at 94°C for 60 seconds then 12 cycles of denaturation at 94°C for 60 seconds, hybridization at 65°C for 60 seconds and elongation at 72°C for 60 seconds; a final elongation was performed for 10 minutes at 72°C. The PCR 2 products were purified using an Agencout AMPure XP bead:DNA ratio of 1:1.25 to obtain libraries. TD product purification, library preparation and paired-end sequencing using an Illumina Hiseq 2000 (1 lane) was performed at the GeT-PlaGe core facility (Genome and Transcriptome, Toulouse) using a TruSeq PE Cluster Kit v3 (2×100 bp) and a TruSeq SBS Kit v3.

### Bioinformatic treatment of TD sequencing

The steps of the informatics treatment from the raw sequencing dataset to population binary (1/0) matrices for the presence/absence of TE insertions per individual are described in Figure S2. First (Figure S2-A), the paired-end reads of each individual were quality checked and trimmed using UrQt v. 1.0.17 (Modolo & Lerat 2015) with standard parameters and a *t* quality threshold of 10. The reads pairs were then checked and trimmed for Illumina adapter contamination using cutadapt (Martin 2011). Specific amplification of TE insertions was controlled by checking for the expected 3′ TE sequence on the R1 read using Blat (Kent 2002) with an identity threshold of 0.90. Only reads with an alignment-length/read-length ratio ≥ 0.90 were retained. The R2 reads for which the R1 mate passed this filter were then selected for insertion loci construction after removal of the TD adapter on the 5′ start using cutadapt and removal of reads under 30 bp. Selected reads were separated in each individual according to the TE families for loci construction.

To correct for the inter-individual coverage variations, we performed a sampling of the cleaned reads (Figure S2-B). First, for each TE family, the distribution of the number of reads per individual was generated and individuals with fewer reads than the first decile of the distribution were removed; then, the cleaned reads of the remaining individuals were randomly sampled at the value of the first decile of coverage (this value varies among TE families). For each TE, the sampled reads of each retained individual were clustered together using the CD-HIT-EST program (Li & Godzik 2006) to recover insertion loci (Figure S2-C). In this all-to-all reads comparison, the alignments needed a minimum of 90 percent identity, the shortest sequence needed to be 95% the length of the longest, global identity was used and each read was assigned to its best cluster. In a second step, the reference reads of each locus within an individual, given by CD-HIT-EST, were clustered with all reference reads of all individuals using the same threshold to build the locus catalog, including a list of loci of all individuals and the coverage for each locus in each individual. After this step, the insertion loci that matched known repeats of the Asian tiger mosquito (Goubert *et al*. 2015) were discarded; alignments were performed with Blastn using the default parameters.

Since the quality control removed a substantial number of reads for the construction of the TE insertions catalog, the raw R2 reads (with their TD adapter removed), that could have been discarded in the first attempt were then mapped over the catalog to increase the scoring sensibility (Figure S2-D). Before mapping, the raw R2 reads were also sampled at the first decile of individual coverage (as described previously). At this step, individuals who were removed from at least two TE families for loci construction were definitively removed from the whole analysis. Mapping (Figure S2-E) was performed over all the insertion loci of all TE families in a single run to prevent multiple hits. Blat was used with an identity threshold of 90 percent. Visual inspection of alignment quality over 30 sampled loci per TE family was performed to ensure the quality of scoring. Raw matrices were then filtered out (Figure S2-F) for a minimum insertion frequency of 2.5% among all individuals and aberrant loci with extreme (> 99th centile) coverage and coverage standard deviations were discarded. The final datasets consisted of one matrix per TE family with information for each individual concerning the presence (1) or absence (0) of TE for each of the selected loci.

To check if the sampling procedure would affect our results, the read sampling procedures and subsequent analysis were performed independently 3 times (replicates M1, M2 and M3).

### Genetic analyses and Genomic scan

The population structure analyses were performed independently for each TE family. Principal coordinates analyses (PCoAs) were performed to identify genetic clusters using the ade4 package (Dray & Dufour 2007) of R 3.2.1 (R development core team 2015). The S7 coefficient of Gower and Legendre (1986) was used as a genetic distance because it gives more weight to shared insertions as follows: with a, b, c and d taken from a contingency table such as a = 1/1 (shared presence); b = 1/0; c = 0/1 and d = 0/0 (shared absence); S7 = 2a /(2a+b+c). Shared absences were not used because they do not provide information on the genetic distance between individuals due to the “copy and paste” mode of transposition of the TE used (shared ancestral state). Pairwise populations *F*_ST_ were computed using Arlequin 3.5 (Excoffier & Lischer 2010); the significance of the index was assessed over 1,000 permutations using a significance threshold of 0.05.

The genomic scan was performed in two steps for each of the sampling replicates of each TE. First, Bayescan 2.1 (Foll & Gaggiotti 2008) was used to test for the deviation of each locus from neutrality. Bayescan considers a fission/island model in which all subpopulations are derived from a unique ancestral population. In this model, variance in allele frequencies between subpopulations is expected to be due either to the genetic drift that occurred independently in each subpopulation or to selection that is a locus-specific parameter. The differentiation at each locus in each subpopulation from the ancestral population is thus decomposed into a *β* component (shared by all loci in a subpopulation) related to genetic drift and a *α* component (shared at a locus by all subpopulations) due to selection. Using a Bayesian framework, Bayescan tests for the significance of the *α* component at each locus. Rejection of the neutral model at one locus is conducted using posterior Bayesian probabilities and controlled for multiple testing using a false discovery rate. In addition, Bayescan integrates uncertainty about allele frequency from dominant data such as the TD polymorphism, leaving the inbreeding coefficient (*F*_IS_) to vary between 0 and 1 during the Markov-Chain Monte-Carlo process. Bayescan was used with default values except for the prior odds, which were set to 100 (more compatible with datasets having thousands of loci, see Bayescan manual) and a significance *q*-value threshold of 0.05 to retain outlier loci.

In a second step, only outliers suggesting divergent directional selection between Europe and Vietnam were considered. To identify these, locus-by-locus analyses of molecular variance (AMOVAs) were performed using Arlequin 3.5 for each TE family. The significance of the *F*_CT_ (hierarchical analogue of the *F*_ST_ measuring the extent of differentiation between groups of populations) between Vietnamese and European populations was assessed by performing 10,000 permutations between individuals among populations with a significance threshold of 0.05. For each dataset, Bayescan outliers were cross-referenced with significant *F_CT_* loci as an objective threshold to restrict the number of candidate loci.

### PCR validation and Outlier analyses

Pairs of primers were designed for each outlier locus to be used in standardized conditions. The forward primer was located in the TE end of the concerned family and the reverse primer was set from the outlier locus (Table S2). All primer pairs were first tested on a set of 10 individuals to assess their specificity using a 1/50 dilution of starting DNA from the TD experiment. Validated primers were then used to check the insertion polymorphisms in 47 representative individuals from the 8 populations studied in the TD experiment using 1/50 dilutions of the starting DNA (not all individuals could be used because of DNA limitations). All PCRs were conducted in a final volume of 25 *μ*L using 0.5 *μ*L of diluted DNA, 0.5 *μ*L of each primer (10 *μ*M), 1 *μ*l of dNTPs (10 mM) and 1 U of DreamTaq Polymerase with 1× green buffer (ThermoFisher Scientific). Amplification was performed as follows: denaturation at 94°C for 2 minutes then 34 cycles including denaturation at 94°C for 30 seconds, hybridization at 60°C for 45 seconds and elongation at 72°C for 45 seconds; a final elongation was performed for 10 minutes at 72°C. After a 45 min migration of the PCR product on a 1× electrophoresis agarose gel, CG and MB assessed the insertion polymorphisms independently.

To identify the genomic environment of the outlier loci, their sequences (reference R2 read) were mapped onto the assembled genome of *Ae. albopictus* assembly AaloF1 strain Foshan (Chen *et al*. 2015) using Blastn. Blastn alignments were performed with default parameters and sorted according to their score. Additionally, each alignment was visually inspected for consistency. Outlier loci with multiple identical hits were discarded. To identify genes surrounding the mapped outliers, we used annotations from VectorBase (http://www.vectorbase.org) for the assembly or, if the *Ae. albopictus* gene was missing annotation, we used the annotation of orthologues in closely related species. In addition, we questioned whether genes potentially involved in the diapause pathway, a critical adaptation required in temperate environments, may be associated with outliers. In a previous publication, Poelchau *et al*. (2013a) identified that differentially expressed genes between diapause induced and non-diapause induced samples were significantly enriched in functional categories related to diapause preparation. These functional categories are defined *a priori* and tested for a more significant differential expression than that expected by chance (Poelchau *et al*. 2013a). Because, at the time of this previous publication, the corresponding transcripts were associated to *Ae. aegypti* orthologues, we here mapped the original transcriptome data (annotated eggs and embryo assembly, downloaded at http://www.albopictusexpression.org) onto the AaloF1 genome assembly using blat with default parameters. After alignment, one best hit was retained per transcript according to the best alignment score. When a transcript had identical best hits, all positions for the transcript were considered. After alignments, the transcript positions were intersected with the AaloF1 gene set using bedtools v2.25.0 (Quinlan & Hall 2010) to identify corresponding *Ae. albopictus* genes. In addition, Chen *et al*. (2015) reported 71 complete diapause-related genes that were merged to our initial candidate set. The eventual enrichment in outliers near these diapause-related genes was assessed using the following procedure: we estimated the total number of base pairs covered by diapause-related genes on the genome assembly and added the longest distance that we found between one outlier and its closer diapause related gene (up to the contig size) to their 5′ and 3′; this defined the “diapause base-pairs”. We then compared the ratio of the number of outliers found within these “diapause base pairs” over the total number of mapped outliers and the ratio of “diapause base-pairs” over the total assembly size using an exact binomial test.

## Results

### High-throughput TE insertion genotyping

The presence/absence of insertions of five TE families were genotyped in an initial number of 140 individuals using a combination of family-specific PCR and individually labelled high-throughput sequencing. Sequencing produced a total of 102,319,300 paired-end reads (2×101 bp). After quality and specificity filtering, 24,332,715 reads were suitable for analyses. The loss of reads was in the great majority due to specificity filtering since quality only resulted in trimming. After application of the read sampling procedure to control for coverage variation between individuals, an average of 128,491 polymorphic insertion loci were available for each of the three sampling replicates. A final 120 individuals were retained per TE family (discarded individuals vary per TE family). The mean number of loci per individual and per TE family ranged from 1025 ± 290 s.d. (IL1 family, mean and s.d. averaged over the three replicates) to 3266 ± 766 s.d. (RTE5 family). Details are given in Table S1. Although our read sampling procedure could have artificially lowered the mean insertion frequency of the loci, this effect should be small because in our final datasets the TE insertion frequencies (*i.e*., the number of individuals who share an insertion) are not influenced by the mean number of reads per individual at the considered locus (Figure S3).

### Population structure

Principal coordinates analyses (PcoAs) were performed independently for each of the five TEs (Figure 2). The shared absence of a specific insertion was not considered in the distance matrices in the PCoAs: Class I retrotransposons have a “copy and paste” transposition mechanism that allows us to infer the “absence” state as the ancestral allele. In addition, these TE insertions segregate at very low frequencies among individuals and thus a shared “absence” is likely to be non-informative with regard to an individual's co-ancestry. Among the three main principal coordinates (PCs), individuals tend to be grouped according to their respective populations with little overlap between groups. However, the three main PCs represent only a small fraction of the total variance (< 10%), suggesting a weak genetic structuring between the populations. Overall, individuals from Vietnamese populations (HCM, TA, VT) tend to be grouped together in a single cluster, with the exception of 13 to 14 individuals from HCM for the L2B and RTE5 TE families (S2 Figure) and six individuals of VT with the RTE4 TE family (Figure 1) that cannot be clearly distinguished from European samples. BCN individuals (Spain) represent the most homogeneous group, well differentiated from Vietnamese and French individuals (SP, CGN, NCE and PLV).

**Figure 2.**
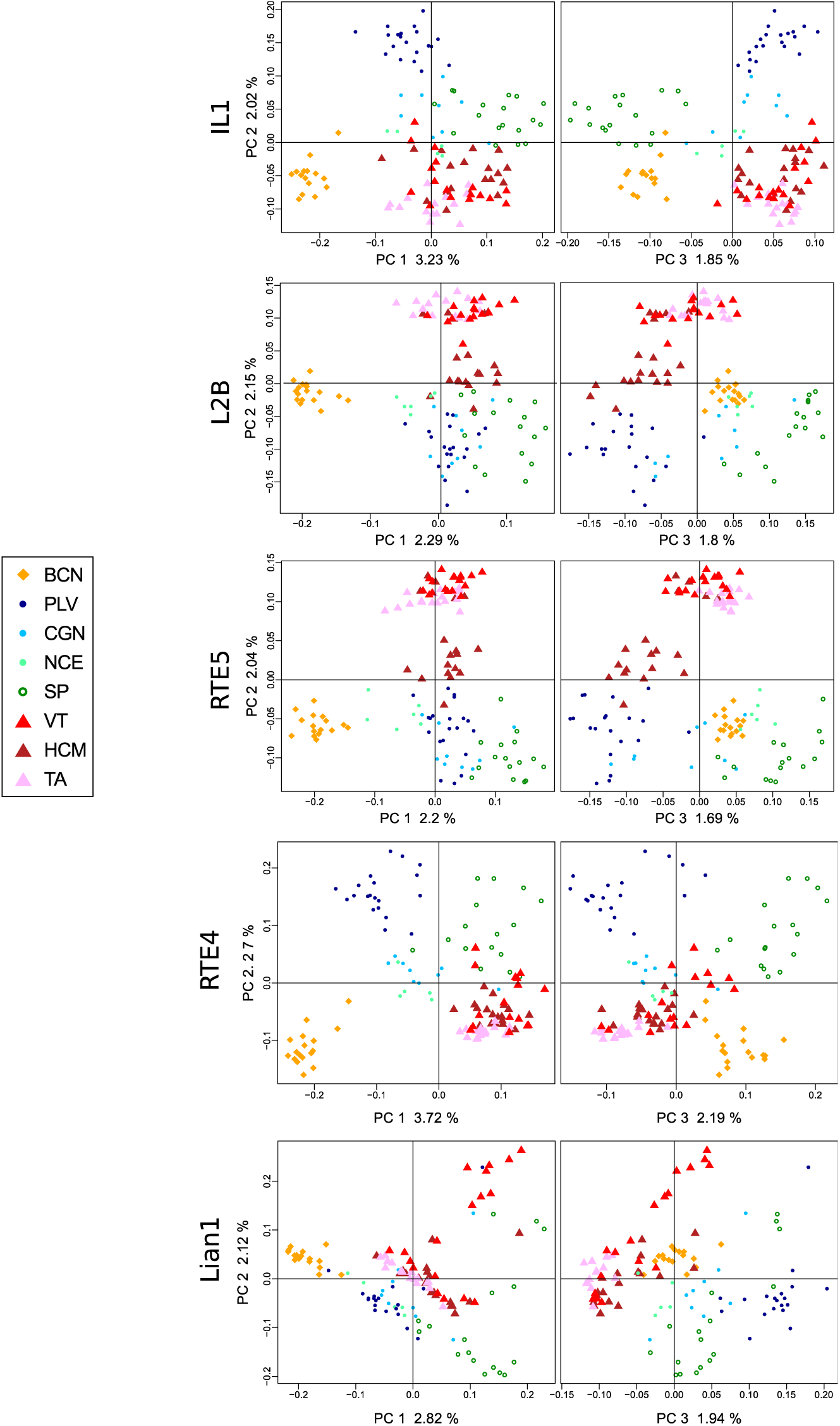
**Principal coordinates analyses (PCoAs).** Projection of individuals over the three first principal coordinates (PC) of PCoAs for each of the 5 TE families and for the first replicate (M1, see Materials and Methods). The proportion of inertia represented by each axes is noted in %. Circles: European populations; triangles: Vietnamese populations. The results for other sampling replicates can be found in Figure S4.

In agreement with the PCoAs, the analyses of molecular variance (AMOVAs) attributed little genetic variance among groups (Vietnam-Europe) and between populations within groups (Table 1). In the studied populations, most of the genetic variance was distributed among individuals within groups.

**Table 1.** Analyses of molecular variance (AMOVAs). The results for the three replicates (M1, M2, M3) of read sampling for the five TE families (IL1, L2B, RTE5, RTE4, Lian1). Values are given in percentage of the total genetic variance.

|  | IL1 | L2B | RTE5 | RTE4 | Lian1 |
| --- | --- | --- | --- | --- | --- |
| Among groups | [0.59-0.70] | [1.22-1.29] | [1.08-1.10] | [1.97-2.04] | [0.67-0.74] |
| Among populations within groups | [5.15-5.37] | [3.58-3.63] | [3.36-3.40] | [6.67-6.78] | [4.47-4.56] |
| Within populations | [94.04-94.16] | [95.08-95.18] | [95.51-95.55] | [91.18-91.30] | [94.77-94.81] |

Intervals report min and max values among the 3 sampling replicates

The measures of genetic differentiation among pairs of populations were consistent between the PCoAs and AMOVAs (File S1): the BCN population shows the highest *F*_ST_ with the other populations for each of the five TEs (0.051 < *F*_ST_ < 0.148) whereas Vietnamese populations were the most closely related (0.011 < *F*_ST_ < 0.032). Although VT is located 100 km away from TA and HCM (both sampled in the same city, Hô Chi Minh, Vietnam), the *F*_ST_ values are very similar between the three Vietnamese populations, suggesting no influence of geography at this scale. CGN and NCE, sampled in the same urban area (Nice agglomeration), also differentiated little or were not significantly differentiated, depending on the TE family. The previously identified intermediate pattern of HCM with some European populations at the L2B and RTE5 loci (PCoAs) is also found at the *F*_ST_ level, especially for the low differentiation with the PLV population for these markers (0.011 < *F*_ST_ < 0.020).

### Genomic scan

Outlier loci for selection signature were searched using Bayescan (non-hierarchical island model) and then sorted for a significant *F_CT_* (between Europe-Vietnam group differentiation) to retain only candidate loci compatible with a differential selection between continents. To reduce false positive risks due to uneven mutation rates between TE families, outlier scans were also performed independently for each TE family (Narum & Hess 2011; de Villemereuil *et al*. 2014). We identified 92 candidate insertion loci (Figure 3). Most of these insertions are found in both areas (no private allele), except for RTE4_1638 and RTE4_1898 that were not found in Vietnam. In addition, 74% of the outliers correspond to high-frequency insertions in Europe, which is significantly more than expected for a 50-50 chance (Chi-squared test, X=20.098, *P* < 0.01) whereas this 50-50 pattern is observed comparing 92 randomly chosen loci with the same overall insertion frequency (≥ 20 individuals/locus, Chi-squared test, X=1.837, *P* = 0.175) between Europe and Vietnam (Figure 4).

**Figure 3.**
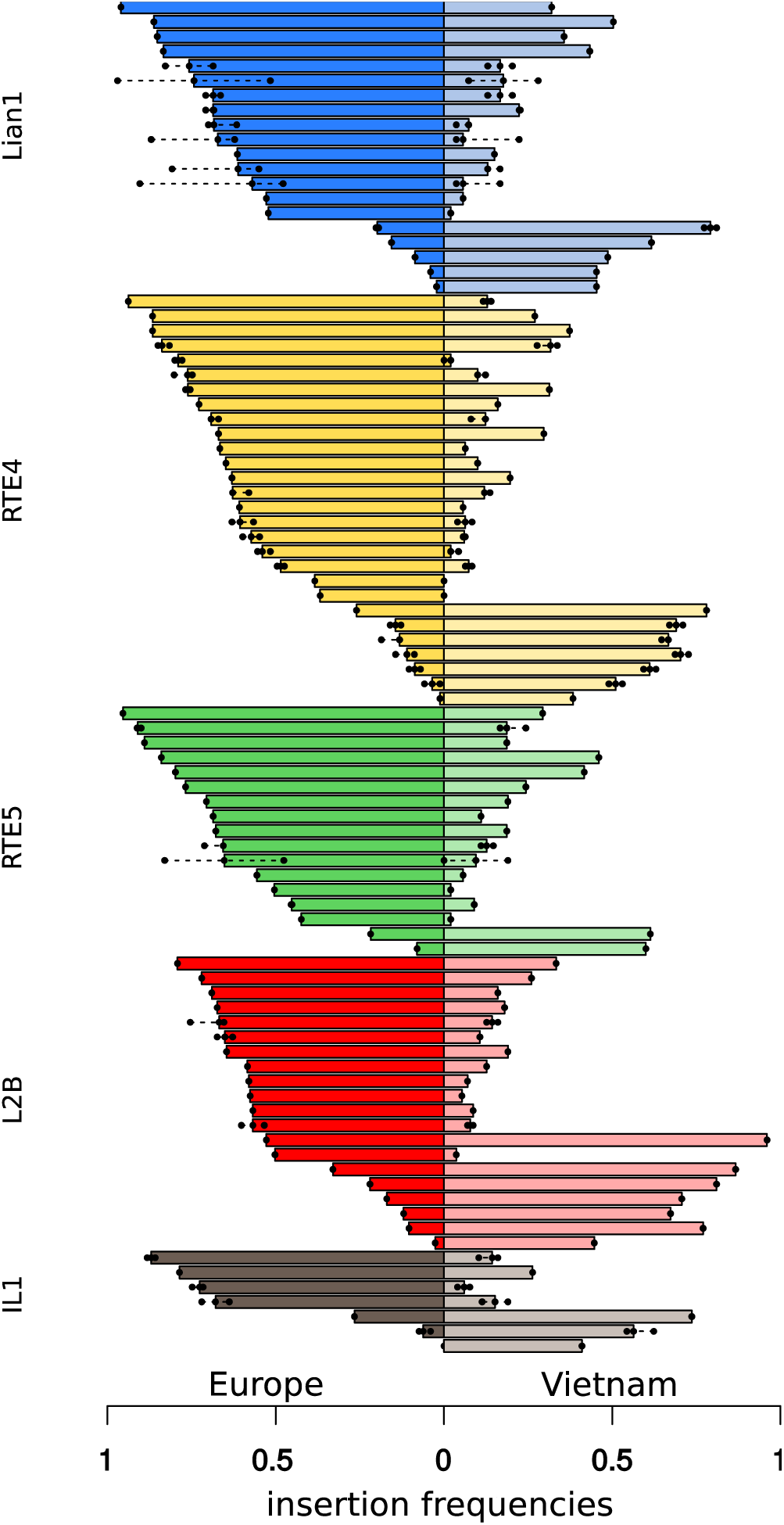
**Insertion frequencies in Europe and Vietnam for 92 outlier loci**. Bars represent the median value from the three reads sampling replicates and dots represent the values from the other replicates (if outlier(s) found in replicates). Colors correspond to each of the 5 TE families.

**Figure 4.**
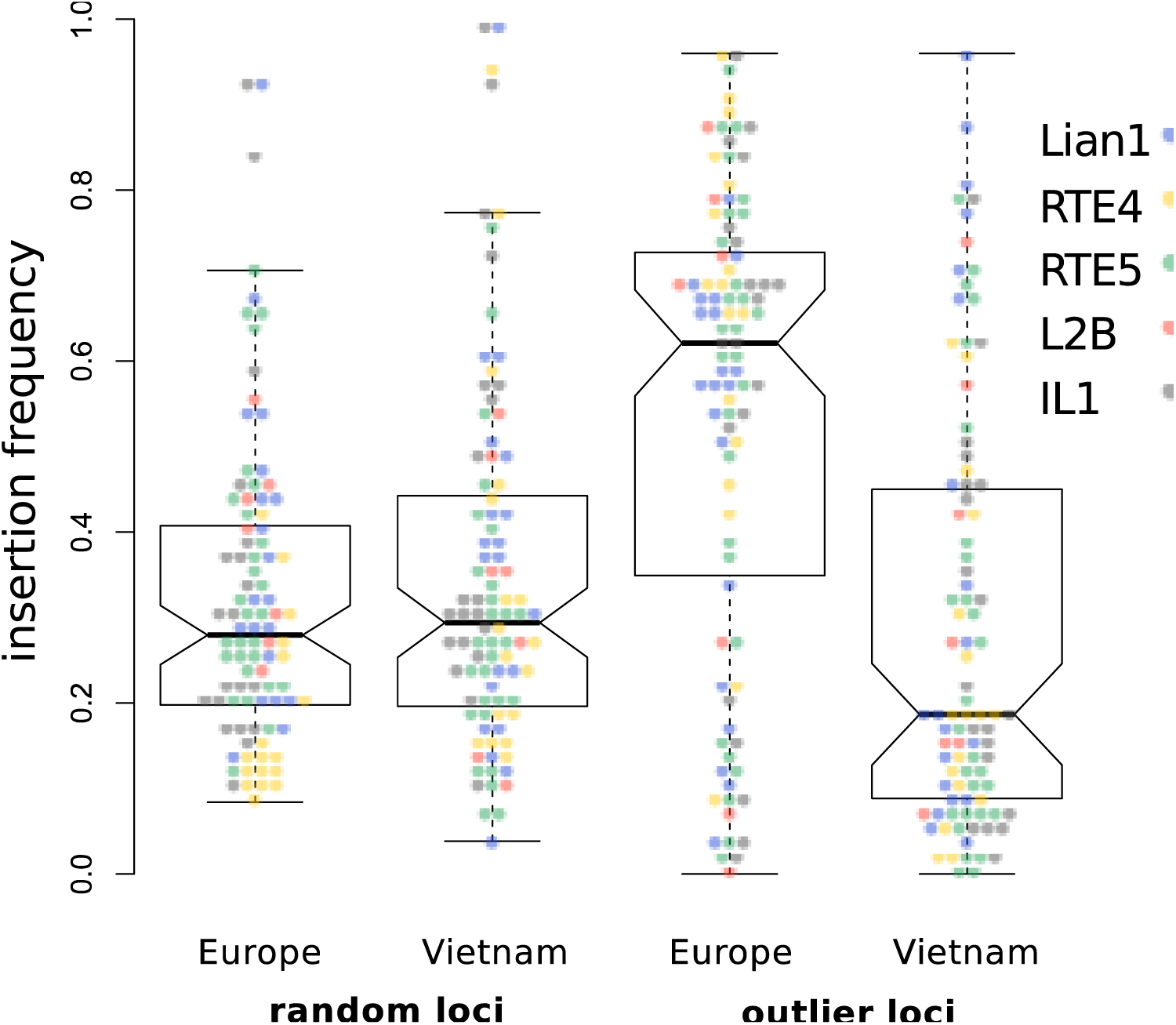
**Comparison of outlier frequencies with randomly selected loci.** Insertion frequencies of 92 randomly chosen loci among those with the same minimum insertion frequency (≥20 individuals) as outliers compared with 92 outlier loci. The random loci were taken from the first replicate (M1) and values for outliers are median values obtained from the three replicates. Non-overlapping notches indicate a significant difference between the true medians (dark horizontal bars).

PCR amplification of the outlier loci was carried out on a representative panel of 47 individuals to validate the insertion pattern detected using TD (see Materials and Method). For loci where the amplification was successful, the insertion pattern observed using PCR always confirmed the TD pattern (Figure S5).

From 92 outlier loci, 21 could be attributed to a unique position on the *Ae. albopictus* genome. Annotation and distance to surrounding genes are reported in the S2 File. Fifteen outliers mapped within contigs with identified genes. We found that 4 outliers (S2 File, sheet 2, highlighted) loci are located on contigs that harbor diapause-related genes. Two of them (Lian1_5902 and RTE4_17015) are located in the direct vicinity, either inside or within 5654 bp, of these genes, which is significantly closer than expected by chance (exact binomial test, *P* = 0.014). Lian1_5902 is located in an intron of *lac1* (longevity assurance factor 1; AALF000670) and RTE4_17015 neighbors the AALF004790 a lipophorin-coding gene. Both genes are known to be involved in lipid metabolic pathways. Although the other diapause related genes are not the closest genes of the two other outliers, they represent two singular groups of genes located in tandem: AALF020842 and AALF00843 located 71.05 kb from RTE5_10123 and AALF020959, AALF020960, AALF020961, AALF020962, AALF020963 and AALF020965 located 216.6 kb from Lian1_11252.

Three other outliers are located within other genes (outliers Lian1_10005, Lian1_9293 and RTE4_34941), including a hemolymph juvenile hormone-binding protein (RTE4_34941/AALF012643). The 10 remaining outliers were located 21.1 kb to 85.2 kb from their closest gene.

## Discussion

The goal of our study was to identify genomic regions involved in adaptive evolution of *Ae. albopictus* thanks to the development of new genetic markers. Through high-throughput genotyping of the insertion polymorphisms of five TE families, we identified up to 128,617 polymorphic loci among more than a hundred of individuals from eight sampling sites. The estimated genome size of *Ae. albopictus* exceeds one billion base-pairs (Goubert *et al*. 2015; Dritsou *et al*. 2015; Chen *et al*. 2015). Accordingly, the number of markers scored in this study offers a comfortable genomic density of one marker every 10 kb.

TE-based methods have been successfully used to perform population genetic analyses within a repetitive genomic environment, such as in the human genome (Watkins *et al*. 2003; Witherspoon *et al*. 2013; Rishishwar *et al*. 2015). Similar high-throughput genotyping methods have been developed for a large panel of organisms (Witherspoon *et al*. 2010; Iskow *et al*. 2010; Sabot *et al*. 2011; Bridier-Nahmias *et al*. 2015; Monden & Tahara 2015) but relied on well-established reference genomes (human, rice, strawberry, yeast). Monden *et al*. (2014) recently completed such an analyses without a reference genome, to score 2024 loci from two TE families in sweet potato. Because of the number of available loci, and being the first of its kind in animals, our study represents a large improvement. We provide a cost-efficient method to quickly generate many polymorphic markers without extensive knowledge of a species' genome. Specifically, this strategy appears extremely valuable for species with a large genome size in which the TE density could severely compromise the development of more classical approaches such as the very popular RAD sequencing.

The genetic structure of the studied populations showed strong consistency between sampling replicates of individual's reads, demonstrating the robustness of the method despite an initial substantial coverage variation among individuals. Population genetics analyses revealed a very low level of genetic structuring between European and Vietnamese populations. Among the studied populations, AMOVAs showed that most of the genetic variation is distributed between individuals within populations (> 90%), and as suggested by pairwise *F_ST_* and PCoAs, only a small part (< 10%) of the genetic variance is due to differentiation between populations. The genetic differentiation we measured is as high among European populations as it is between populations from Europe and Vietnam.

This singular population structure is in agreement with previous results gathered in *Ae. albopictus* using different collections of allozymes, mtDNA or microsatellite markers (Black *et al*. 1988; Kambhampati *et al*. 1991; Zhong *et al*. 2013; Gupta & Preet 2014; Manni *et al*. 2015). Moreover, a recent analysis performed with a set of 11 microsatellites on individuals from the same populations (with the exception of BCN) showed a similar distribution of genetic variation among hierarchical levels (Minard *et al*. 2015). These results demonstrate the reliability of our markers and confirm that a non-hierarchical island model can likely fit the global genetic structure. The genetic diversity observed in Europe is compatible with a scenario of multiple and independent introductions, as already suggested for *Ae. albopictus* (Urbanelli *et al*. 2000; Birungi & Munstermann 2002; Takumi *et al*. 2009; Becker *et al*. 2013). However, as previously suggested, this pattern could also be the result of founder events that may occur during colonization and/or a restriction of gene flow between populations after their introduction. Answering such a question would require extended sampling over the entire native area.

The outlier analysis revealed 92 loci with high posterior probabilities of being under positive selection between European and Vietnamese populations. When possible, the PCR amplification of the outlier loci using a set of representative individuals confirmed a shift of insertion frequencies toward either the European or the Vietnamese sampling sites. This suggests that, despite reduced coverage, introduced by sampling in the dataset, the scored insertion polymorphisms are reliable. In addition, our method of analysis is likely to be conservative: indeed, the Bayescan outliers were selected for their consistency with a significant *F_CT_* between European temperate and Vietnamese tropical populations, which avoids retaining outliers that we were not looking for, for example those due to a population-specific event.

Interestingly, we found significantly more outlier loci with a high frequency in Europe and low frequency in Vietnam. This was unexpected under our initial assumptions as follows: a favored allele selected in one or another environment has *a priori* no reason to be more often associated with the presence or the absence of a TE insertion at linked sites. A possible explanation is that the majority of the sequenced TE insertions segregate at low frequencies (approximately 10% of all individuals). When considering the linked region of one polymorphic TE insertion, if a favorable mutation appears in an individual in which the insertion is absent, the increase of frequency of this “absence” haplotype will thus, most of the time, have a modest effect on the genetic differentiation at this marker because it is already segregating at high frequency. In contrast, if a favorable mutation appears in a TE “presence” haplotype, the increase in frequency of the linked TE insertion would lead to high differentiation (*F_CT_*). In absence of an alternative explanation, our outlier loci could thus indicate in which subset of populations the adaptive mutation occurred and in the present case this would have occurred more frequently in the temperate populations. Additionally, the observation of such a low global insertion frequency is expected to reduce the risk of false positive outliers due to negative selection acting on potentially deleterious polymorphic insertions.

Two scenarios that are not mutually exclusive could be invoked in light of our data. A simple case would be a direct adaptive evolution in European invasive population that originated from tropical regions of the native area. A second hypothesis could be that invasive temperate populations came from the northernmost territories of the native area such as northern China or Japan where *Ae. albopictus* populations are already cold-adapted. It would be thus interesting to know whether the observed signature of selection results from more “ancient” adaptations in the native area or if it originates from more recent fine tuning of cold-related traits in the invasive areas. A recent study (Porretta *et al*. 2012) using new variable *COI* mtDNA sequences and historical species range modeling suggested that northern territories of the native area of *Ae. albopictus* would be the latest to have been colonized after a range expansion from Southern refugia after the last glacial approximately 21,000 years ago. The authors suggested that *Ae. albopictus* may have followed the human populations during their expansion from South to North in this area that began approximately 15,000 years ago. Thus, regardless of the origin of the invasive individuals sampled in Europe, it is likely that they are representatives of populations that had recently undergone a shift of selective pressure from tropical to temperate climatic conditions. This could explain why so many outliers are associated with high insertion frequency in Europe and why some candidate genes in the diapause pathway are found in the neighborhood of some of these outliers. An easy method to distinguish between these possibilities would be to search if the same outlier insertions are present in several temperate populations from the native area.

We were able to assign a unique position for 21 of the outlier loci on the *Ae. albopictus* genome. Sixteen outliers were found inside or in the close vicinity of annotated genes, which may allow speculation on potential targets of selection. These genes encompass functions related to cell structure and organization, lipid metabolism, or signal transduction. A main challenge faced by temperate populations is overwintering (Mori *et al*. 1981; Hawley 1988; Takumi *et al*. 2009; Denlinger & Armbruster 2016). Previous studies have shown that the cell cycle and lipid metabolism are specifically solicited at multiple stage of the diapause preparation and maintenance (Urbanski *et al*. 2010; Poelchau *et al*. 2013a; b; Huang *et al*. 2015) to allow temperate populations to go through winter within cold-resistant eggs. Although the genes found in our genome scan have not been functionally associated with diapause, two of them *laf1* (AALF000670) and a lipophorin-coding (AALF004790) are a differentially expressed between induced and non-induced samples during diapause preparation (Poelchau et al., 2013a) and are located closer than expected by chance from two of our outlier loci. These genes are both involved in lipid metabolism and could be thus strong candidates for adaptive evolution. Another notable candidate is AALF012643, a hemolymphatic carrier of the juvenile hormone (JH). JH appears to be critical in the maintenance of the diapause *in Ae. albopictus* but its exact function remains to be elucidated (Poelchau *et al*. 2013b). Although appropriate caution should be taken regarding the sole candidate status of these genes, it is worth mentioning that the diapause pathway has already been shown to benefit from rapid adjustments due to local adaptation in *Ae. albopictus*. For instance, Urbanski *et al*. (2012) showed that invasive American populations originating from Japan have rapidly evolved a new adaptive clinal response to diapause induction, independent from that observed in the native area. Thus, adaption in the temperate regions may have led to several selective sweeps on gene or regulatory sequences involved in this critical pathway, allowing the settlement of the mosquito in new temperate areas. Further experiments, including fine-scale study of the genetic diversity of these candidates among populations, are needed to assess their potential implication in the adaptation of *Ae. albopictus* toward temperate environments. Specifically, targeted resequencing of the candidate regions, including outliers, genes and their flanking regions across several individuals and populations should help to determine evidence of selective sweep, the precise extent and location of such events, and eventually designate the causative selected mutations.

It is also important to note that the results presented here are only restricted to a subset of the Asian tiger mosquito populations located in temperate and tropical environments. Thus, it is probable that some of the outliers detected could be specific to this particular comparison and do not reflect the global pattern of differentiation between tropical and temperate populations. Research on the same outliers between other tropical and temperate populations from native and non-native areas would be extremely valuable to extrapolate our results at a larger scale and refute possible false positive. Should the same outlier insertions be found at high frequencies in temperate locations – such as in USA, Japan or China –, extended investigations about the origin of invasive populations would help clarify if those similarities are due to an ancestral sweep or parallel sweeps that occurred independently in several populations. This study already provides a set of functional primers for some candidate loci that could be directly used to answer this question in *Ae. albopictus* DNA samples. As with every novel method, our study may be susceptible to unforeseen or underrated bias; though we attempted to remove these issues (such as controlling the relationship between loci coverage and insertion frequency), we identified several points that must be taken into account for interpretation of the results. First, unlike random SNPs, TE insertions can be potential targets of purifying selection, which can sometimes mimic the diversity pattern of a selective sweep (Charlesworth *et al*. 1993; Stephan 2010). However, as reported earlier, insertions are found at low frequencies and negative selection for the “presence” of TE is likely to produce a slight change in the allele frequency spectrum. Another issue may be the uncertainty related to the null model used in the genome scan; even if we did not explicitly evaluate the fit of our data to a specific population model, we did not find evidence to reject an island model such as that implemented in Bayescan. This software is also the best suited to handle our dominant data but is indeed restricted to mainly detect recent, strong and monogenic positive selection (Pérez-Figueroa *et al*. 2010; Narum & Hess 2011; de Villemereuil *et al*. 2014); additionally, it should be insightful to compare our results with more diverse genome-scan models and we would like to emphasize that our results were produced under the specific hypothesis of the model used.

Here, we report the first information supporting adaptive evolution at the molecular level in the Asian tiger mosquito. Progress in the annotation of published genomes and the looming availability of supplementary genomic resources will allow the most gain from these results. We hope that this work will contribute to unraveling the implications of adaptive processes during the invasion of disease vectors.

## Acknowledgements

We thank Van Tran-Van, Christophe Bellet, Grégory Lambert, Huynh Kim Ly Khanh and Trang Huynh who made possible and contributed to sampling in France and Vietnam. We are grateful to Valèria Romero Soriano and her family for their help during sampling in Sant Cugat dèl Vallès. Library construction and sequencing was performed in collaboration with Clémence Genthon and Olivier Bouchez. We thank Manon Vigneron for PCR validation experiment work. We also thank Rita Rebollo, who provided insightful comments and English revision of the manuscript. This work was performed using the computing facilities of the CC LBBE/PRABI. C.G. received a grant from the French Ministry of Superior Education. This work was supported by the Centre National de la Recherche Scientifique and the Institut Universitaire de France and preliminary experiments benefited from a grant from the Federation de Recherche 41 “Bio-Environnement et Santé”. Funding for mosquito sampling in Vietnam was provided by grants from EC2CO CNRS and occurred within the framework of GDRI “Biodiversity and Infectious Diseases in Southeast Asia”. Original maps used to describe sampling were downloaded from http://d-maps.com/carte.php?num_car=4719&lang=en and http://d-maps.com/carte.php?num_car=708&lang=en.

## Data Accessibility

Paired-end raw sequences are available through SRA at NCBI under SRP070185 (Bioproject PRJNA312147)

Final presence/absence matrices (including replicates) are available at Dryad (doi:10.5061/dryad.9p925) at http://datadryad.org/review?doi=doi:10.5061/dryad.9p925

## Author contributions

CG, CV and MB conceived of the experiments and conducted the analyses. CG and HH developed and performed the molecular experiments. GM, CVM and PM conducted the sampling in France and Vietnam. All authors contributed to the final version of the manuscript.

